# Distinct Neural Signatures Underlie Finger Tapping and Walking to Auditory Rhythms

**DOI:** 10.64898/2026.07.30.741770

**Authors:** C. Ziane, D. Schön, S. Dalla Bella

**Affiliations:** International Laboratory for Brain, Music and Sound Research (BRAMS), University of Montreal, Pavillon Marie-Victorin, 90 Vincent D’Indy Ave Local A-108, Montréal, Québec H2V 2S9, Canada; School of Kinesiology and Physical Activity Sciences, University of Montreal, Centre d’éducation physique et des sports, 2100 Édouard-Montpetit Blvd, Montréal, Québec H3T 1J4, Canada; Department of Psychology, University of Montreal, Pavillon Marie-Victorin, 90 Vincent D’Indy Ave, Montréal, Québec H2V 2S9, Canada; Centre for Research on Brain, Language and Music (CRBLM), 2001 McGill College Ave, Montréal, Québec H3A 1G1, Canada; VIZJA University, Okopowa 59, Warsaw, 01-043, Poland; Institut de Neurosciences des Systèmes (INS), Inserm, Aix-Marseille University, 27 Jean Moulin Blvd, Marseille, 13005, France; Institute of Language, Communication and the Brain (ILCB), CNRS, Aix-Marseille University, 5 Pasteur Ave, Aix-en-Provence, 13604, France

## Abstract

Neural oscillations synchronize with rhythmic stimuli, shaping perception and action. Auditory-motor synchronization (AMS) enhances this process but most evidence comes from finger tapping, an attentionally demanding voluntary behaviour. Whether motor-driven modulation of neural synchronization generalizes to more automatic movements like walking remains unclear. In this study, we used mobile EEG to compare neural synchronization to auditory rhythms during tapping and walking in 40 older male and female participants, manipulating cognitive load (single *vs*. dual task) and task instructions (synchronize with *vs*. ignore metronome). EEG components attuned to metronome frequencies were extracted and complementary measures of neural synchronization were computed (phase coupling to the stimulus, power at the stimulus frequency, and the stability of instantaneous frequencies). Both movements enhanced neural synchronization relative to passive listening. Neural synchronization was further increased by instructions to synchronize, but only during tapping. This instruction-facilitating effect was lost in dual tasks. These findings suggest that AMS recruits movement-dependent neural and cognitive mechanisms, depending on where AMS lies on a continuum from voluntary-controlled to automatic forms of coordination, with implications for rhythm-based interventions in aging.

**Significance statement:** Neural coupling to stimuli in the environment shape perception and action, but its dependence on movement type and cognitive control is poorly understood. Using mobile EEG, we show that both finger tapping and walking enhance neural coupling to the auditory stimuli, but tapping is uniquely modulated by task factors. Our findings suggest that cortical involvement during rhythmic motor tasks differs for voluntary (tapping) and automatic (walking) behaviours. As gait can move from being mostly automatic to voluntary-controlled (*e.g.*, under heightened cognitive load, in motor disorders), it offers a unique opportunity to study the dynamic interactions between sensory, motor, and cognitive processes supporting neural coupling to auditory stimuli.

## Introduction

Rhythmic auditory stimuli can synchronize neural activity and behaviour across multiple temporal scales (Harding et al., 2025). This phenomenon, often referred to as *entrainment*, is thought to reflect the alignment of endogenous neural oscillations with external rhythmic input (Lakatos et al., 2019). By aligning periods of high neural excitability with predictable sensory events, entrainment is proposed to support temporal prediction, attentional selection, and sensorimotor coordination (Lakatos et al., 2019). Neural entrainment to auditory rhythms has been consistently observed using electroencephalography (EEG), notably through increases in oscillatory power at the stimulation frequency and phase coupling to the stimulus (Henry & Obleser, 2012; Nozaradan et al., 2011). Importantly, auditory rhythms not only shape neural dynamics during passive listening, but also strongly influence motor behaviour (Rosso et al., 2023).

Neural entrainment can be facilitated by auditory-motor synchronization (AMS), the ability to coordinate movements with rhythmic sounds. For example, synchronizing finger taps to metronomes increases neural entrainment relative to passive listening (De Pretto et al., 2018; Nozaradan et al., 2015). Previous studies have also shown that neural entrainment can be modulated by attentional demands and task instructions, indicating that this process is not only stimulus-driven but also shaped by cognitive and motor processes (Haegens & Zion Golumbic, 2018). Importantly, AMS can emerge spontaneously, without explicit instructions (Zagala et al., 2024) or be voluntary, likely relying more on attentional and executive control (Leow et al., 2014). Together, these findings suggest that neural entrainment reflects dynamic interactions between sensory, motor, and cognitive systems.

Numerous studies investigated AMS through finger-tapping tasks (Repp, 2005). Finger tapping provides a controlled and experimentally viable model for laboratory research on AMS. However, it is a relatively artificial behaviour that depends strongly on voluntary control and attentional resources (Ziane et al., 2026). Findings from tapping may thus not generalize to other more widespread and natural rhythmic behaviours, such as gait. Walking is intrinsically periodic and partially reliant on spinal structures which afford automatic control (Dimitrijevic et al., 1998). In everyday contexts, humans spontaneously adapt their gait to external stimuli. Voluntary adjustments are achieved through cortical control (Delval et al., 2020). AMS during gait can be both spontaneous and voluntary depending on individual traits (*e.g*., rhythmic abilities, cognitive functions) (Leow et al., 2018; Zagala et al., 2024). AMS during tapping and walking may thus rely on partially distinct neural and cognitive mechanisms (Ziane & Dalla Bella, 2025).

While finger-tapping paradigms dominate AMS research, neural mechanisms supporting gait, an ecologically valid, partially automatic behaviour, remain poorly understood, largely due to substantial movement-related artifacts in EEG signals (Castermans et al., 2014). To our knowledge, only one study investigated neural entrainment during gait (Chen et al., 2025), but it lacked synchronization to auditory rhythms, a critical premise for motor-driven facilitation of auditory processing (Morillon et al., 2014). Here, we directly compare neural entrainment during tapping and walking to auditory rhythms matching preferred movement frequencies.

To probe automatic *vs*. voluntary mechanisms, we manipulated cognitive load and task instructions. We recently proposed that rhythmic behaviours lie along an automatic-voluntary continuum (Ziane et al., 2026). For example, gait is mostly automatic in healthy individuals. Reliance on voluntary control can however be achieved by increasing cognitive load with a secondary task. When gait shifts toward voluntary control, timing performance becomes linked to finger-tapping performance. We manipulated cognitive load using single and dual tasks and directed attention toward and away from auditory stimuli through task instructions. Neural entrainment was quantified using complementary measures capturing phase coupling to the stimulus, power at the stimulus frequency, and instantaneous frequencies. We first hypothesized that movement would enhance neural entrainment relative to passive listening. More importantly, we predicted that neural entrainment would be more strongly modulated by task instructions and cognitive load during tapping than walking based on the idea that gait is more automatic. Finally, we examined whether neural entrainment predicted behavioural synchronization performance across movement types and task factors.

## Methods

Methods described below were part of a larger project, from which behavioural data has already been published in a preprint article (Ziane et al., 2026). Information described in some of the sections below thus matches the method section of that study.

### 1. Participants

Forty older adults (26 females), aged 69.88 ± 3.91 (mean ± SD), with diverse educational backgrounds (17.09 ± 3.56 total years of education), with and without musical training volunteered to participate in the study. Most participants (78 %) had received some form of informal music training throughout their lives (*e.g.*, music classes offered in the regular school curriculum) but only four participants had received seven or more years of formal training (*e.g.*, private lessons, conservatory). All participants were right-handed (thus right-footed; (Peters & Durding, 1979)), and cognitively intact as assessed by a score ≥ 26/30 on the Mini-Mental State Examination (Folstein et al., 1975; Mitchell, 2017). Exclusion criteria included having any auditory, motor, psychological or neurological condition, and use of medication affecting the nervous system. All participants provided informed consent to participate in this study, which was approved by the University of Montreal’s Ethics committee for research in education and psychology (#2024-5766).

### 2. Experimental design

Participants completed two laboratory sessions within one week (**Figure 1**).

**Figure 1.**
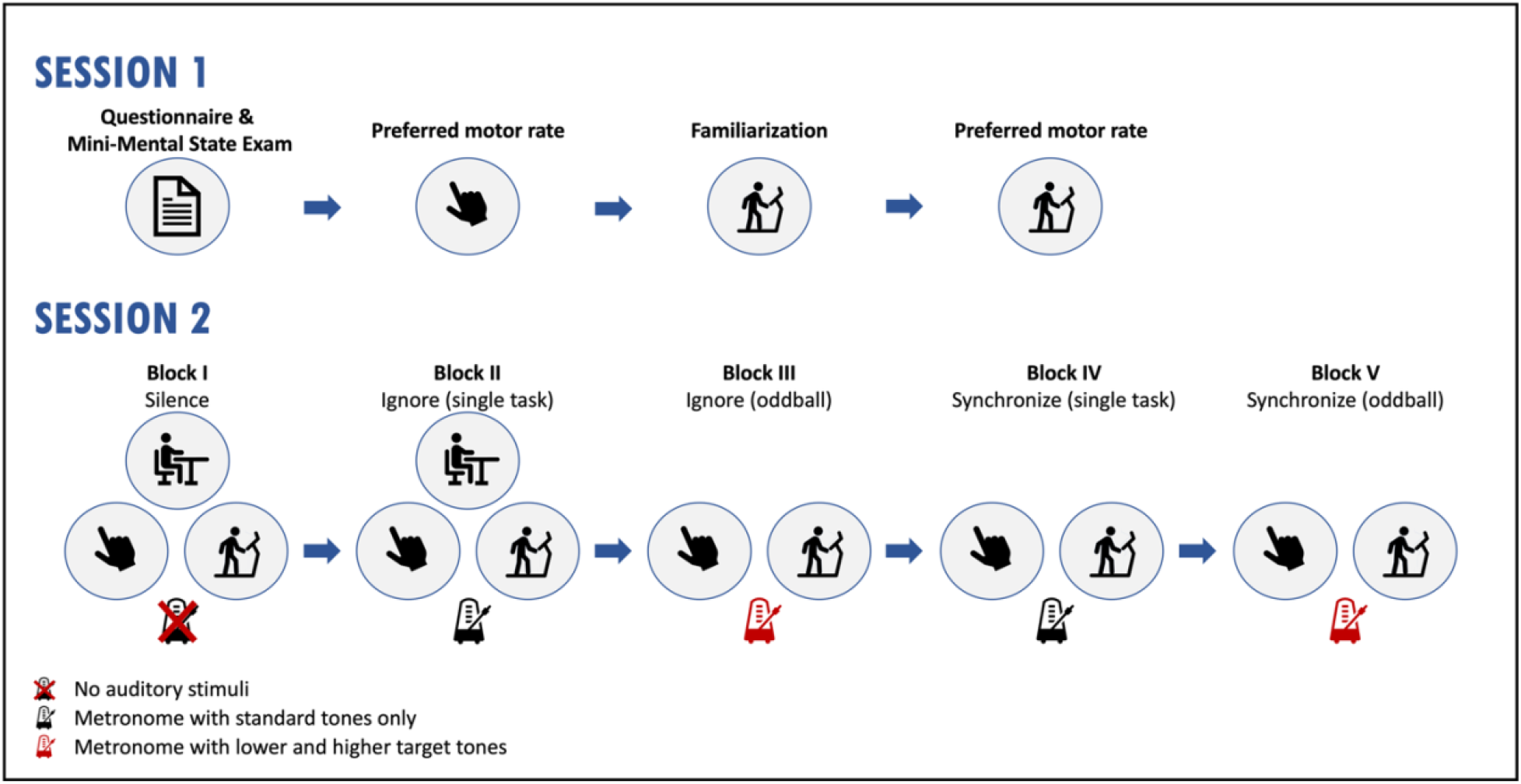
Experimental design. During the Session 1, participants completed a questionnaire about their demographic information and musical expertise. Then, we assessed their cognitive state with the Mini-Mental State Examination, and their preferred motor rates during tapping and walking. During Session 2, participants were equipped with mobile EEG before completing twelve 5-minute conditions. Blocks I to V were completed in a predetermined order, while conditions within blocks were randomized.

#### 2.1. Session 1: Calibration and familiarization

During Session 1, we collected demographic information, assessed musical expertise and assessed cognitive status. Participants’ preferred motor rates for tapping and walking were measured, and they were familiarized with the experimental tasks.

##### Tapping rate

Participants were instructed to tap as regularly as possible for one minute at a comfortable, self-selected pace.

#### Walking rate

After treadmill familiarization, we estimated the participant’s walking speed via an adaptive procedure. Treadmill speed was set to 2.6 km/h and increased by 0.2 km/h increments until participants reported reaching their preferred speed. The treadmill was then restarted at a speed 5 % faster and participants readjusted the speed until their preferred walking pace was achieved again. This procedure was repeated until the participants decreased the speed of the treadmill. Preferred walking speed was defined as the average of the ascending and descending self-selected speeds. Participants then walked at this speed for one minute to determine step rate. Preferred speeds ranged from 2.40 to 5.15 km/h (3.92 ± 0.65 km/h).

##### Oddball familiarization

To ensure participants could discriminate target from standard tones and walk safely on the treadmill while completing the oddball task, we created an audio file containing standard and target tones presented at an inter-beat interval (IBI) matching preferred motor rate. Participants walked for one minute while counting the number of high and low target tones and reported both numbers at the end.

#### 2.2. Session 2: Experimental task

During Session 2, participants were equipped with EEG and completed twelve 5-minute conditions involving resting, finger tapping, and walking to a metronome matching preferred motor rate. Experimental conditions were organized into five blocks with different instructions (ignore or synchronize to the stimuli) and cognitive loads (single or dual task). In dual tasks, participants performed an auditory oddball concurrently with movement.

In Block I (silence), participants performed each movement without auditory stimulation. In Blocks II-III (ignore), participants were instructed to maintain their spontaneous movement rates while ignoring auditory stimuli. In Blocks IV-V (synchronize), participants were instructed to synchronize their movements with auditory stimuli (*i.e.*, voluntary synchronization). Within each block, conditions were presented in randomized order. Block order was fixed to minimize carry-over effects related to instruction and stimulus exposure. In oddball blocks (blocks III and V), participants were told that performance on the cognitive and the motor task was as important.

In single-task conditions, the auditory stimulus consisted of a metronome sequence of pure tones at 600 Hz presented at an IBI matching each participant’s preferred motor rates.

In dual-task oddball conditions, ∼100 standard tones (regardless of total number of tones) were replaced with low (300 Hz) and high frequency (1200 Hz) target tones. Target tones were pseudo-randomly distributed, with a minimum of two standard tones between successive targets.

#### 3. Instrumentation

*EEG.* Cortical activity was recorded with a 64-channel mobile EEG system (LiveAmp, Brain Vision, Munich, Germany), following the 10-20 system for electrode placement. Impedances were kept below 25 kΩ. Signals were recorded at a sampling frequency of 500 Hz.

##### Finger taps

A force signal was recorded using a custom-made device, equipped with a force sensitive resistor (FSR).

##### Walking steps

Vertical forces were recorded using an instrumented treadmill equipped with two force platforms (Tandem treadmill, AMTI, Watertown, USA).

##### Audio delivery

Stimuli were delivered to the participants binaurally with insert earphones (ER-2 Tubephone insert earphones, Lucid Hearing Holding Company, Fort Worth, USA) at a comfortable level.

Movement and audio data were integrated and synchronized in the Qualisys Track Manager software (Qualisys, Göteborg, Sweden) and recorded at 2000 Hz. Data from all systems were synchronized by sending a TTL pulse from Matlab to both the Qualisys and the EEG systems and aligning recordings offline.

#### 4. Data processing

All data processing was done in Matlab 2021b with custom-made scripts, unless otherwise specified. Scripts are available on Github (https://github.com/claraziane/projetDT).

#### 4.1. EEG data

Signals were processed with the BeMoBIL pipeline (Klug et al., 2022), developed for mobile brain/body imaging data. Line noise at 60 Hz was removed with the Zapline-plus plugin (Klug & Kloosterman, 2022). Signals were re-referenced to the average to detect bad channels based on correlations with surrounding electrodes (correlation threshold: .7). Detected bad channels were interpolated using spherical spline interpolation from unreferenced signals and referenced. Adaptive mixture independent component analysis (AMICA) was performed on the data high-pass filtered at 1.5 Hz. The resulting decomposition was applied to the broadband data. Only components flagged as brain-related by ICLabel (Pion-Tonachini et al., 2019) were kept. Clean signals were high-pass filtered at 0.2 Hz. Final data quality was verified through visual inspection in both temporal and frequency domains. This procedure let to the rejection of 5.8 % of the data (2 ± 2 participant/condition).

#### 4.2. Extraction of auditory-related components

From the 64-channel signals, we extracted one EEG component that was maximally attuned to the frequency of the auditory stimulus. To do so, we created a spatial filter using generalized eigendecomposition (Cohen & Gulbinaite, 2017; Rosso et al., 2021). Generalized eigendecomposition identifies spatial filters that maximize the contrast between two covariance matrices:

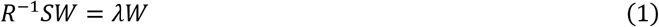

where R is the reference covariance matrix, S is the covariance matrix to be maximized, W is a square matrix that contains as many weight vectors (*i.e.*, eigenvectors) as there are channels, and λ is a diagonal matrix, which contains the eigenvalues (*i.e.*, contrast ratios) associated with each eigenvector. R was computed from broadband EEG signals (Moumdjian et al., 2025; Rosso et al., 2021), while S was computed from the EEG signals gaussian-filtered around the auditory frequency (Full-Width at Half Maximum = .3 Hz) (Cohen & Gulbinaite, 2017; Rosso et al., 2021). S and R were computed for each stimulus (-100 to +500 ms around onset). In silent conditions (block I), the stimulus frequency and onsets of the corresponding condition in block II were used to compute S and R. Euclidean distances from mean S and R were computed. Averaged S and R were recomputed after removal of matrices with z-scores > 2.23, to exclude particularly noisy segments (Rosso et al., 2021). Generalized eigendecomposition was performed separately for each condition.

The five components best separating S and R were obtained from the eigenvectors associated with the largest eigenvalues. Topographies and frequency spectra were inspected. Components were selected based on (i) a clear spectral peak at the stimulus frequency and (ii) a physiologically plausible topography. No component could be extracted for one participant in two experimental conditions. From extracted components, we computed phase coupling to the stimulus, power at the stimulus frequency, and the stability index. Note that when there was no stimulus (block I), stimulus frequency (equivalent to preferred movement rate) and onsets of the corresponding condition in block II were used to compute phase coupling, power and the stability index.

##### Phase coupling

The component time series was band-pass filtered around the stimulus frequency. Then, we applied a Hilbert transform to get instantaneous phases. Phase values at stimulus onsets were used to compute the resultant vector length, reflecting phase consistency across stimuli (Berens, 2023). Values were logit transformed prior to statistical analysis.

##### Power at the stimulus frequency

Power spectra were computed using a fast Fourier transform. To remove the 1/f structure, power was expressed as signal-to-noise ratio (SNR) (Cohen & Gulbinaite, 2017), defined as power at the stimulus frequency relative to neighbouring frequencies (+/− .5 Hz, excluding +/− .1 Hz). SNR values were summed across adjacent bins to account for spectral leakage (+/− .02 Hz).

##### Stability index

Instantaneous frequencies were derived from the unwrapped phase angles (Cohen, 2014). A moving median filter (window width: 400 frames, order: 10) was then applied to attenuate nonphysiological fluctuations. The stability index was computed as the standard deviation of instantaneous frequencies, with lower values indicating more stable tracking of the stimulus (Rosso et al., 2021).

#### 4.3. Behavioural data

Tap and step onsets were extracted from force signals. Synchronization performance was quantified using synchronization consistency (Dalla Bella et al., 2024; Sowiński & Dalla Bella, 2013). Asynchronies between movement and stimulus onsets were converted into phase angles, and the resultant vector length was computed (Berens, 2023).

### 5. Statistical analyses

First, we ensured participants had synchronized their movements to the auditory stimuli by computing Rayleigh’s test on phase angles, for each condition and participant. A significant Rayleigh’s test indicates that phase values are not uniformly distributed around a circle (*i.e.*, the participant synchronized).

Then, two linear mixed models were fitted separately for each neural measure (phase coupling, power, stability index) after outlier rejection (|z-scores| ≥ 3). Partial eta squared 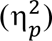 were computed to assess effect sizes.

#### Auditory-entrainment model

Only rest conditions from blocks I and II, and tapping/walking conditions from blocks I and IV were used. We were only interested in comparing rest and tapping, as well as rest and walking with and without auditory stimulation. The following equation was used:

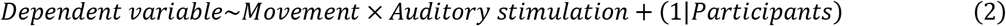

#### Task-factor model

Only movement conditions with auditory stimuli were used. The following equation was used to assess main effects of movement type (2 levels: tapping, walking), cognitive load (2 levels: single task, oddball), and instruction (2 levels: ignore, synchronize).

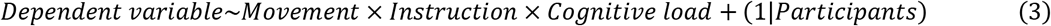

Finally, to assess brain-behaviour relationships, Spearman correlations were computed between neural measures and synchronization consistency after outlier removal. Normality was assessed prior to analysis.

All *p*-values were Bonferroni-corrected, with α = 0.05. Statistical analyses were conducted with RStudio 4.3.0.

## Results

### 1. Participants synchronize their movements with auditory rhythms regardless of instruction

Participants synchronized their movements to the auditory stimuli in most conditions (**Figure 2**). Under voluntary synchronization (instructions to synchronize), 38/40 participants synchronized during tapping (2 failed in the single task; 3 in the oddball task), and 39/40 synchronized during walking (1 failed in the single task; 3 in the oddball task) (Rayleigh test, *p* < .05).

**Figure 2.**
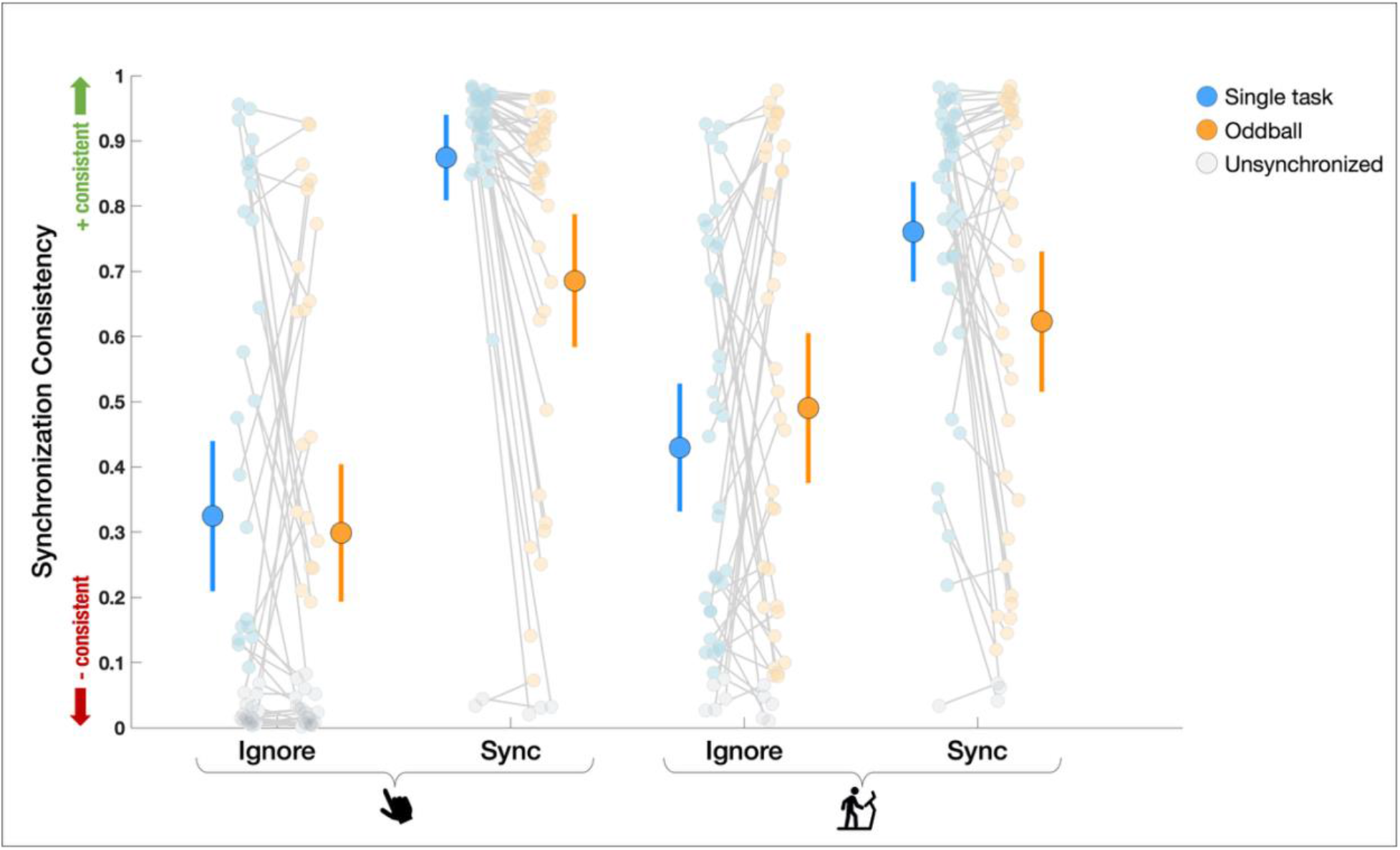
Synchronization consistency. Group mean with 95 % confidence-interval bars and participants’ individual values in single-task (blue) and oddball (orange) conditions under ignore and synchronization instructions. Participants who did not synchronize (p_Rayleigh_ ≥ .05) are shown in grey.

Even when instructed to ignore the metronome, most participants synchronized their movements, although more robustly for walking than tapping. Under ignore instructions, 35/40 participants synchronized during walking, compared to 23/40 and 21/40 in single-task and dual-task tapping, respectively^1^.

### 2. Movement enhances neural entrainment to auditory rhythms

To rule out movement artifacts as a source of entrainment-like signals, we compared rest, tapping, and walking in the absence of auditory stimulation (**Figure 3**). If neural entrainment was primarily driven by the production of rhythmic movements, tapping and walking should exhibit stronger entrainment than when at rest. Contrary to this prediction, without auditory stimuli, no differences in phase coupling and power emerged between rest and movements (phase coupling, tapping *vs* rest: *t*(179) = 0.09, *p* = 1.00, walking *vs* rest: *t*(180) = 1.80, *p* = 0.148; power, tapping *vs* rest: *t*(175) = 0.48, *p* = 1.00, walking *vs* rest: *t*(178) = 2.01, *p* = 0.092), confirming that rhythmic movement alone does not generate entrainment-like neural signatures. The stability index did not differ between rest and tapping (*t*(177) = -1.69, *p* = 0.187) but was lower during walking than rest (*t*(177) = -2.78, *p* = 0.012), suggesting gait may involve endogenous rhythmic neural dynamics.

**Figure 3.**
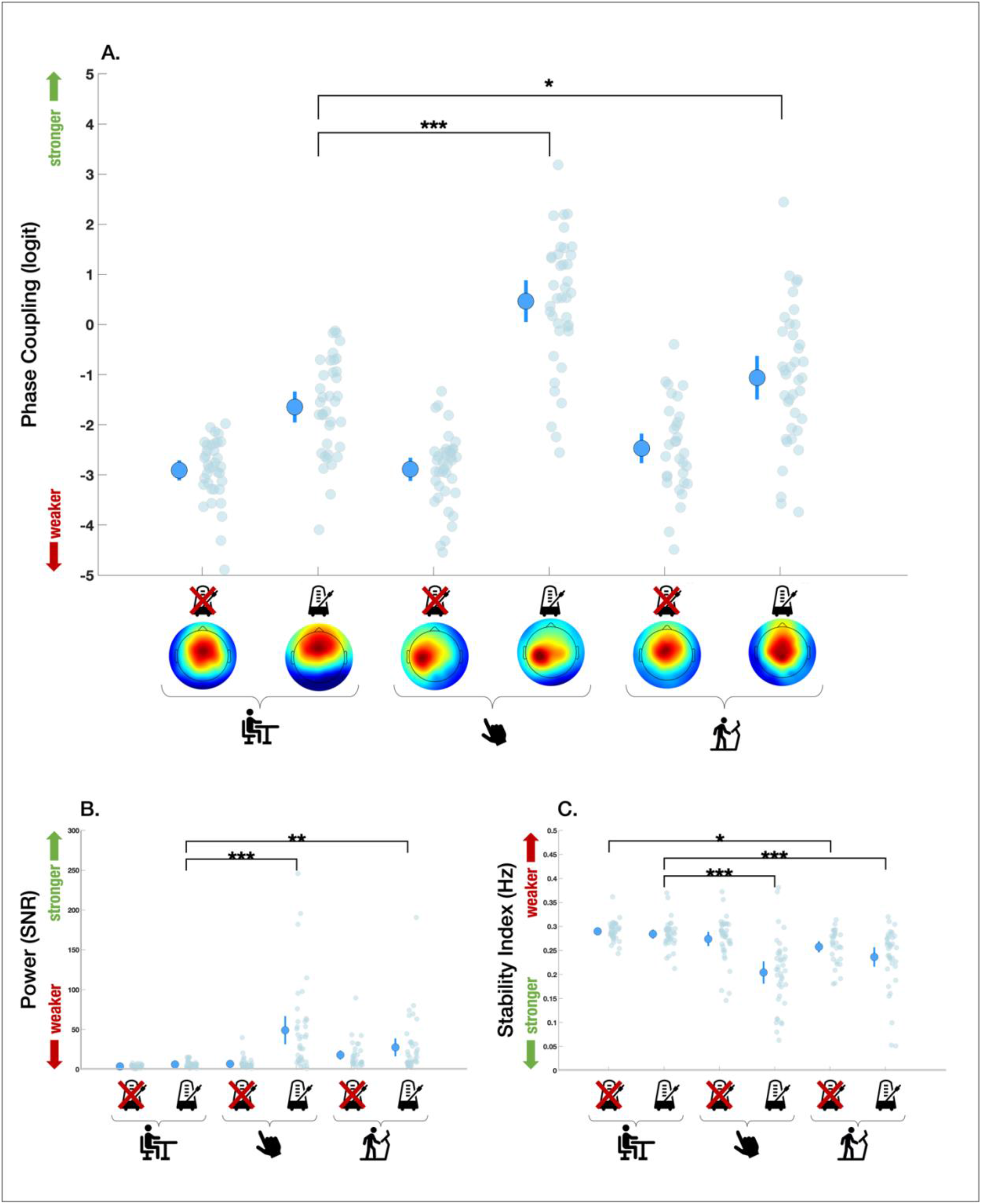
Results from auditory-entrainment model. Group mean with 95 % confidence-interval bars and participants’ individual **A.** phase coupling to the beat, **B.** power at the beat frequency, and **C.** stability indexes. Significant differences are indicated with single, double, and triple asterisks for *p* ≤ .05, *p* ≤ .01, and *p* ≤ .001, respectively.

In the presence of auditory rhythms, movement amplified neural entrainment across all measures. Compared to rest, phase coupling and power were higher during tapping (phase coupling: *t*(178) = 9.23, *p* < .001; power: *t*(173) = 6.56, *p* < .001) and walking (phase coupling: *t*(177) = 2.53, *p* = .025; power: *t*(174) = 3.27, *p* = .003). Similarly, the stability index was lower (more stable tracking) during tapping (*t*(176) = -7.70, *p* < .001) and walking (*t*(175) = -4.54, *p* < .001).

Together, these results indicate that neural entrainment arises from auditory-motor interactions, and not merely from movement alone.

### 3. Task demands differentially modulate neural entrainment during tapping and walking

Having established that neural entrainment is enhanced during movement only in the presence of auditory stimulation, we next examined how it is modulated by task demands, namely movement type, instruction, and cognitive load.

#### 3.1. Voluntary synchronization selectively enhances entrainment during tapping

We first tested whether instructions to synchronize with the auditory stimulus would enhance neural entrainment relative to ignoring it, and whether this effect depended on the type of movement. If top-down control facilitates entrainment, effects should be stronger during voluntary synchronization.

Voluntary synchronization increased entrainment during tapping across all measures (phase coupling: *t*(260) = -7.73, *p* < .001; power: *t*(254) = -4.97, *p* < .001; stability index: *t*(257) = 5.40, *p* < .001). In contrast, no effect of instruction was observed during walking.

Furthermore, under voluntary synchronization, entrainment was greater for tapping than walking, as reflected in phase coupling (*t*(261) = 7.39, *p* < .001) and the stability index (*t*(257) = -2.69, *p* = .031), but not power (*t*(257) = 2.32, *p* = .085). When ignoring the stimulus, entrainment was comparable across movements, although power showed a marginal increase during walking (*t*(256) = -2.51, *p* = .051)^2^.

Overall, these results indicate that instructions to synchronize selectively enhance neural entrainment during tapping, with limited effects during walking, suggesting tapping relies more on top-down control.

#### 3.2. Cognitive load attenuates entrainment under voluntary synchronization

Having shown that instructions selectively enhance entrainment for tapping, we next tested whether cognitive load (dual-task oddball) would further modulate this effect. If entrainment relies on attentional resources, dual-task conditions should weaken it, particularly when participants are voluntarily synchronizing.

Dual tasking reduced entrainment only under voluntary synchronization (phase coupling: *t*(261) = 3.19, *p* = .007; power: *t*(253) = 3.34, *p* = .004; stability index: *t*(257) = -2.73, *p* = .027)^3^. In contrast, no cognitive load effect was observed in ignore conditions.

In single-task conditions, synchronization instructions increased entrainment (relative to ignoring the stimulus) across all measures (phase coupling: *t*(260) = -6.69, *p* < .001; power: *t*(252) = - 4.58, *p* < .001; stability index: *t*(256) = 4.64, *p* < .001). When dual tasking, synchronization instruction still enhanced entrainment, but only for phase coupling (*t*(261) = -2.63, *p* = .036)^4^.

Overall, these findings indicate that facilitation of entrainment through instructions to synchronize is lost under high cognitive load. Top-down modulation of entrainment in a synchronization task thus depends on available cognitive resources.

### 4. Neural entrainment predicts behavioural synchronization depending on attention and instructions

Finally, we investigated whether neural entrainment was functionally related to behavioural synchronization performance. If this is the case, stronger entrainment should be associated with more consistent alignment of movements to the beat.

In line with this prediction, we observed robust brain-behaviour relationships during tapping when participants ignored the auditory stimulus. Stronger neural entrainment was associated with improved synchronization performance across all measures: phase coupling (single task: *ρ*(37) = .75, *p* < .001; oddball: *ρ*(36) = .80, *p* < .001; **Figure 5A**.), power (single task: *ρ*(35) = .70, *p* < .001; oddball: *ρ*(35) = .62, *p* < .001; **Figure 5B**.), and the stability index (single task: *ρ*(36) = -.46, *p* = .033; oddball: *ρ*(35) = -.49, *p* = .018; **Figure 5C**.)^5,6^.

**Figure 4.**
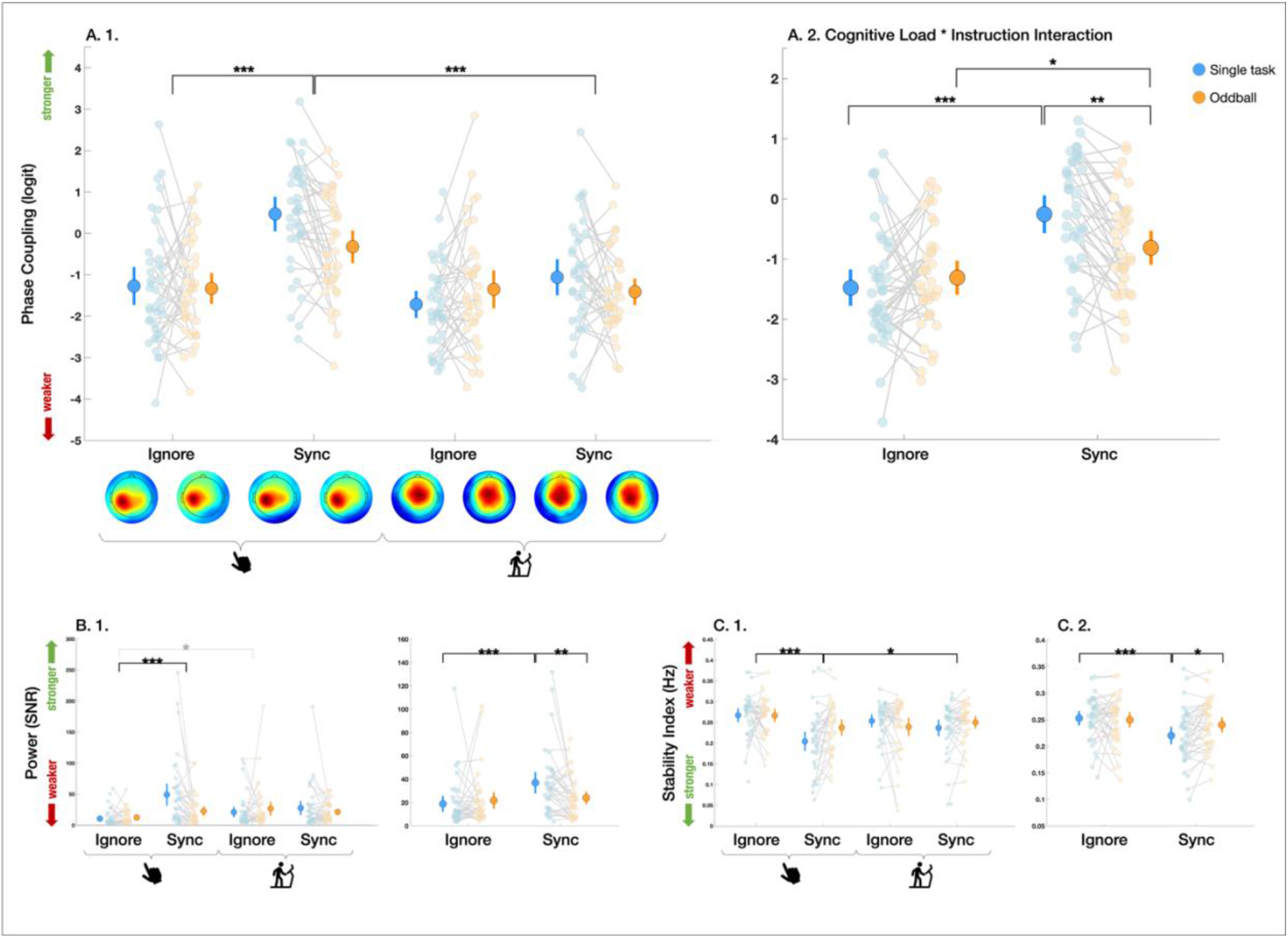
Results from task-factor model. Group mean with 95 % confidence-interval bars and participants’ **A.** phase coupling to the stimulus, **B.** power at the stimulus frequency, and **C.** stability indexes. Single-task conditions appear in blue, while oddball conditions are in orange. In **A.2.**, **B.2.**, and **C.2.**, results are averaged over movements to show cognitive load × instruction interactions. Significant differences are indicated with single, double, and triple asterisks for *p* ≤ .05, *p* ≤ .01, and *p* ≤ .001, respectively. Marginally significant differences are in grey.

**Figure 5.**
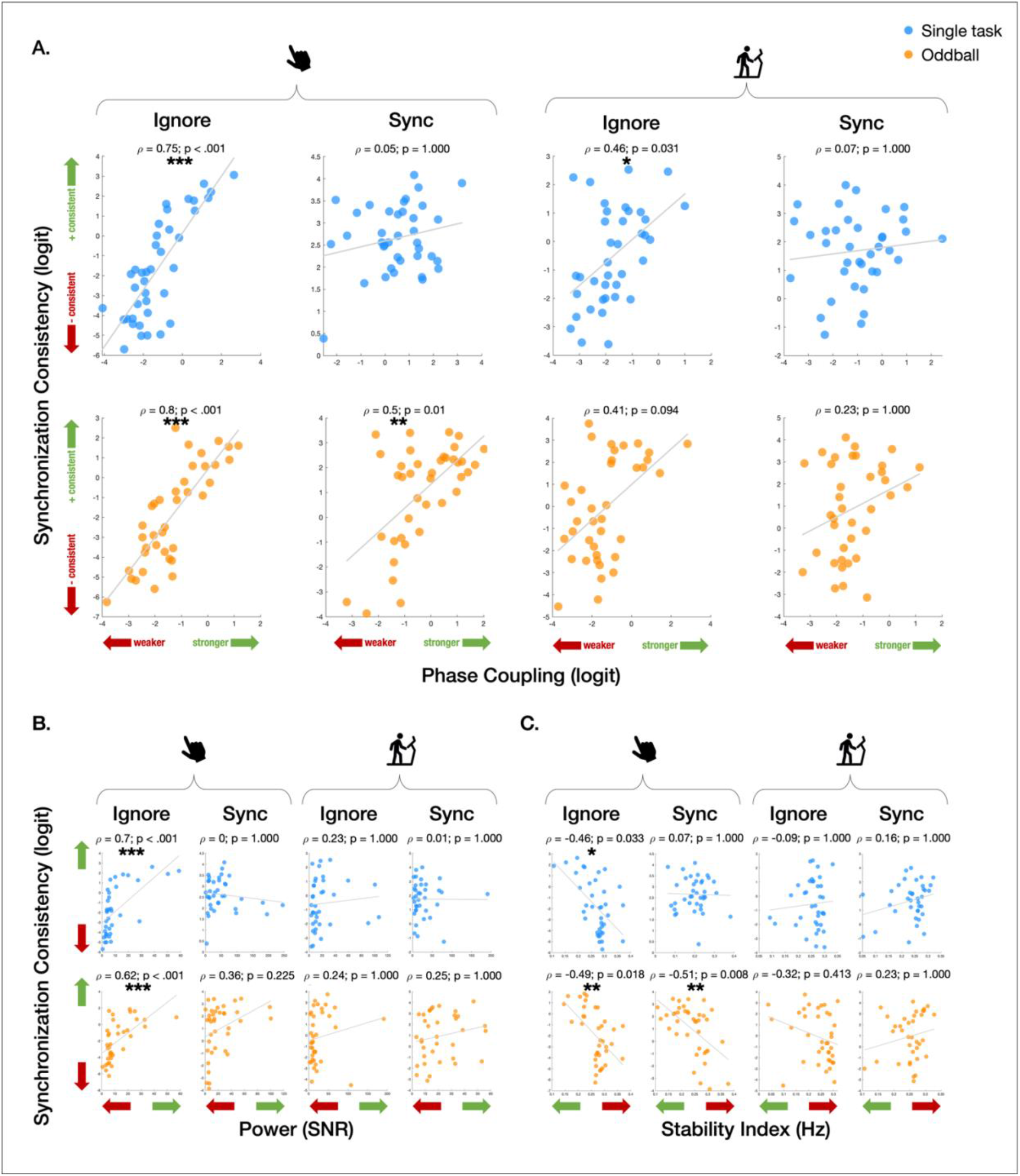
Correlation analyses. Relationships between synchronization consistency and **A.** phase coupling to the beat, **B.** power at the beat frequency, and **C.** stability indexes in single-task (blue) and oddball (orange) conditions for tapping (left) and walking (right) under ignore and synchronization instructions. Spearman’s rho and p-values are indicated for each condition. Significant differences are indicated with single, double, and triple asterisks for *p* ≤ .05, *p* ≤ .01, and *p* ≤ .001, respectively. Regression lines are for visualization purposes only.

When instructed to synchronize and under high cognitive load, brain-behaviour relationships were more selective. In this condition, tapping performance was associated with phase coupling (*ρ*(37) = .50, *p* = .010) and the stability index (*ρ*(37) = -.51, *p* = .008), but not power (*ρ*(36) = .36, *p* = .225)^7^.

Lastly, a more limited relationship was observed during walking: phase coupling was positively associated with synchronization performance in the single-task ignore condition (*ρ*(36) = .46, *p* = .031)^8^, but no consistent associations were found across other conditions.

Overall, neural entrainment is generally associated with better behavioural synchronization in ignore conditions, particularly for tapping. Under voluntary synchronization, relationships are weaker and task-dependent, suggesting that neural entrainment may more directly support behaviour under automatic rather than voluntary AMS.

## Discussion

Neural entrainment was captured during AMS tasks. Auditory stimulation elicited robust neural entrainment, further enhanced by movement (*vs*. passive listening), both for tapping and walking. Notably, instructions modulated neural entrainment differently for tapping and walking: instructions to synchronize enhanced entrainment during tapping, whereas walking remained unaffected. Cognitive load abolished this facilitation, suggesting that top-down control of entrainment requires attentional resources. These findings support a continuum of AMS control, from a voluntary (tapping) to a more automatic (walking) form of synchronization. Considering the challenges associated with mobile EEG (Castermans et al., 2014), reliably capturing neural entrainment during gait tasks was a technical achievement.

A major concern with mobile EEG is movement-related artefacts at frequencies (Castermans et al., 2014), overlapping the auditory beat. While gait’s periodicity alone could produce entrainment-like signals, silence conditions showed no such increase: phase coupling and power did not differ between rest and motor tasks without auditory stimulation. The lack of increase in entrainment by movement in absence of auditory stimuli suggests that our preprocessing pipeline (BeMoBIL, AMICA, ICLabel) and component-extraction method successfully minimized movement-related artefacts, enabling reliable assessment of entrainment to auditory rhythms during gait.

Our first hypothesis was that movement would enhance neural entrainment relative to passive listening, consistent with previous AMS studies (De Pretto et al., 2018; Nozaradan et al., 2015). In line with this prediction, tapping and walking both increased neural entrainment relative to rest. As discussed above, motor-facilitation of entrainment was not observed in silence conditions. These findings support the idea that auditory rhythms provide a stable temporal scaffold that organizes neural activity, while movement enhances the alignment between endogenous neural dynamics and external rhythms. Such facilitation may reflect reciprocal auditory-motor coupling, consistent with the active sensing framework proposing that motor activity contributes to the temporal prediction of sensory events (Morillon et al., 2015; Morillon et al., 2014; Schroeder et al., 2010). Supporting this view, cortical and subcortical motor regions are recruited during rhythm perception (Chen et al., 2008) and rhythmic movements have been shown to enhance auditory processing (Cason et al., 2015; Chen et al., 2025; Morillon et al., 2014). In this context, AMS may emerge through dynamic coupling between externally driven auditory signals and internally generated motor processes.

Interestingly, the stability index was lower during walking than rest in silent conditions, suggesting that locomotor rhythmicity may generate endogenous temporal dynamics even in the absence of external pacing. While these dynamics could reflect internally generated gait rhythms, these effects were substantially weaker than those observed with auditory stimulation, indicating that endogenous motor rhythms alone cannot fully account for the neural signatures associated with synchronization.

Our second hypothesis was that neural entrainment would be more strongly modulated by task instructions and cognitive load during tapping than walking. Consistent with this prediction, instructions to synchronize enhanced entrainment during tapping, but not walking. The effect was lost under heightened cognitive load, suggesting that top-down modulation of entrainment requires attentional resources. Differences between tapping and walking likely reflects partially distinct timing mechanisms. We recently proposed that rhythmic behaviours laid somewhere on an automatic-voluntary continuum (Ziane et al., 2026). AMS performance for tapping and walking only correlate when gait is driven towards voluntary control through heightened cognitive load (Ziane et al., 2026). When gait is mostly automatic when cognitive load is low, timing abilities (motor variability, rate, synchronization consistency) do not correlate with those measured while tapping, suggesting distinct timing mechanisms supporting both behaviours. Neural data from the current study supports the idea that tapping relies more on voluntary control while walking relies more on automatic control (Ziane & Dalla Bella, 2025).

Walking is intrinsically rhythmic and partially supported by spinal and subcortical locomotor mechanisms, including central pattern generators (Dimitrijevic et al., 1998; Gerasimenko et al., 2010; Nielsen, 2003). Although cortical activity contributes to gait regulation under complex conditions, such as obstacle avoidance, direction changes, or irregular terrains (Delval et al., 2020; Seeber et al., 2014; Wagshul et al., 2019), steady treadmill walking is likely to require less voluntary control than finger tapping (Ziane et al., 2026). If AMS during gait is more automatic than during tapping, it is also less sensitive to instructions. Walking showed weaker overall entrainment and fewer brain-behaviour relationships than tapping, further supporting reduced cortical involvement in locomotor synchronization. Future studies should investigate how neural entrainment during gait changes under conditions requiring greater voluntary control.

A secondary objective of this study was to examine whether neural entrainment predicted behavioural synchronization performance across movement types and task factors. Stronger neural entrainment was associated with greater synchronization at the behavioural level, but only in specific conditions. Stronger relationships emerged in ignore conditions, contrasting with previous studies showing links under voluntary synchronization during tapping (Nozaradan et al., 2016; Nozaradan et al., 2015). This discrepancy may reflect lower cortical involvement needed when stimulus tempo matches spontaneous motor rates.

For example, the stability index correlated with synchronization consistency in Rosso et al. (2021). In this study, participants synchronized to a fixed 1.67 Hz tempo, considered to approximate the spontaneous motor rate of the general population (Repp, 2005). However, preferred tapping rates vary tremendously across individuals (Hammerschmidt et al., 2021) and AMS performance is influenced by the relationship between stimulus and spontaneous motor frequencies (Harding et al., 2025). It is thus possible that the 1.67 Hz tempo in Rosso et al. (2021) was far enough from participants’ spontaneous rates, necessitating top-down control to guide behaviour.

When stimulus frequencies match preferred motor rates, high synchronization consistency can be achieved without relying on stimulus information by simply maintaining a stable motor rhythm. Indeed, healthy older adults can produce highly stable motor output during tapping (Dalla Bella et al., 2024), walking (Ready et al., 2022), and even dual-task walking (Ziane et al., 2026). In contrast, tapping becomes more variable under dual-task demands, particularly in older adults (Krampe et al., 2010), increasing reliance on external temporal cues. In these cases, entrained neural oscillations may be particularly useful for guiding behaviour. Thus, restriction of brain-behaviour relationships to ignore conditions could reflect top-down modulation of entrainment to guide motor output. Alternatively, absence of brain-behaviour relationships under voluntary synchronization may be due to a ceiling effect: almost all participants displayed very high AMS during single-task tapping with instructions to synchronize, leaving little variability to correlate with neural measures.

Despite our best effort to reliably extract entrainment measures from EEG components, three limitations should be mentioned. First, neural entrainment measures were computed relative to the auditory stimulus. As a result, the present analyses were specifically designed to capture synchronization to the auditory rhythm rather than internally generated motor rhythms. Endogenous neural dynamics related to self-paced movement may have been present but at different frequencies. Second, despite combining multiple preprocessing and analysis approaches to enhance signal-to-noise ratio, mobile EEG data during walking is vulnerable to residual motion artifacts (Gorjan et al., 2022; Kline et al., 2015; Seeber et al., 2015). Entrainment-like neural signatures however did not systematically emerge during rhythmic movement in silence, arguing against a purely movement-driven explanation of the observed effects. Finally, the present brain-behaviour relationships remain correlational and cannot establish causality. Future studies combining non-invasive brain stimulation or perturbation paradigms may help clarify the causal role of neural oscillations in AMS.

In conclusion, this study demonstrates that neural entrainment to auditory rhythms depends not only on external rhythmic stimulation, but also on movement performed and task factors. Consistent with previous AMS studies, movement enhanced auditory neural entrainment relative to rest. For the first time, we show that walking also increases entrainment of slow neural oscillations to an auditory beat, extending previous findings obtained primarily during finger tapping paradigms. Tapping appeared more sensitive to instructions and attentional demands than gait, which was fairly stable across task factors. These results suggest that different motor behaviours engage partially distinct forms of AMS, from mostly voluntary to automatic forms of synchronization. As gait can move from automatic to voluntary (*e.g.*, heightened cognitive load, motor disorders) forms of synchronization (Ziane et al., 2026), it offers a unique opportunity to study both top-down and bottom-up mechanisms supporting neural entrainment to auditory rhythms. Indeed, our findings support the idea that neural entrainment involve dynamic interactions between sensory, motor, and cognitive systems. In the present work, we highlight the potential of mobile EEG approaches for investigating neural entrainment during ecologically relevant behaviours, opening new perspectives for studying rhythmic coordination in aging and clinical populations.

## Acknowledgments

This research was supported by the Fonds de recherche du Québec - Nature et technologies (FRQNT) doctoral training scholarship, awarded to Clara Ziane; as well as by a Discovery Grant (RGPIN-2019-05453) from the Natural Sciences and Engineering Research Council of Canada (NSERC) and the Canada Research Chair program (CRC in Music Auditory-Motor Skill Learning and New Technologies), awarded to Simone Dalla Bella. The authors would like to thank Louane Bonnemberger, Jeremy Rueda and Océane Martin for their help with data collections, as well as all the participants who took part in this study.

The authors disclose the use of artificial intelligence tools for language editing purposes only.

## Footnotes

1 Note that the following analyses and figures include all participants. Analyses were however repeated without participants who did not synchronize. Footnotes are included whenever this procedure changed the results.

2 This effect appeared for phase coupling (*t*(216) = 3.91, *p* < .001) but disappeared for power (*t*(221) = -1.09, *p* = 1.00) when repeating analyses without participants who did not synchronize.

3 This effect disappeared for the stability index (*t*(205) = -2.43, *p* = .065) when repeating analyses without participants who did not synchronize.

4 This effect disappeared when repeating analyses without participants who did not synchronize (*t*(211) = -.82, *p* = 1.00).

5 This effect disappeared in the single task for the stability index (*ρ*(19) = -.49, *p* = .205) when repeating analyses without participants who did not synchronize.

6 This effect disappeared in the oddball task for phase coupling (*ρ*(17) = .60, *p* = .065), power (*ρ*(16) = .25, *p* = 1.00), and the stability index (*ρ*(16) = -.52, *p* = .227) when repeating analyses without participants who did not synchronize.

7 This effect disappeared for phase coupling (*ρ*(34) = .39, *p* = .142) and the stability index (*ρ*(34) = -.42, *p* = .087) when repeating analyses without participants who did not synchronize.

8 This effect disappeared when repeating analyses without participants who did not synchronize (*ρ*(31) = .34, *p* = .411).

